# GNS561 exhibits potent *in vitro* antiviral activity against SARS-CoV-2 through autophagy inhibition

**DOI:** 10.1101/2020.10.06.327635

**Authors:** Philippe Halfon, Eloïne Bestion, Keivan Zandi, Julien Andreani, Jean-Pierre Baudoin, Bernard La Scola, Jean-Louis Mege, Soraya Mezouar, Raymond F. Schinazi

**Author notes:** **Corresponding author:** Philippe Halfon, M.D, Ph.D., 10, Rue d’Iéna, 13006 Marseille, France, Mail. Contributed equally (Philippe Halfon, Eloïne Bestion, Soraya Mezouar, Raymond F. Schinazi).

## Abstract

Since December 2019, severe acute respiratory syndrome coronavirus 2 (SARS-CoV-2/2019-nCoV) has spread quickly worldwide, with more than 29 million cases and 920,000 deaths. Interestingly, coronaviruses were found to subvert and hijack the autophagic process to allow their viral replication. One of the spotlights had been focused on the autophagy inhibitors as a target mechanism effective in the inhibition of SARS-CoV-2 infection. Consequently, chloroquine (CQ) and hydroxychloroquine (HCQ), a derivative of CQ, was suggested as the first potentially be therapeutic strategies as they are known to be autophagy inhibitors. Then, they were used as therapeutics in SARS-CoV-2 infection along with remdesivir, for which the FDA approved emergency use authorization. Here, we investigated the antiviral activity and associated mechanism of GNS561, a small basic lipophilic molecule inhibitor of late-stage autophagy, against SARS-CoV-2. Our data indicated that GNS561 showed the highest antiviral effect for two SARS-CoV-2 strains compared to CQ and remdesivir. Focusing on the autophagy mechanism, we showed that GNS561, located in LAMP2-positive lysosomes, together with SARS-CoV-2, blocked autophagy by increasing the size of LC3-II spots and the accumulation of autophagic vacuoles in the cytoplasm with the presence of multilamellar bodies characteristic of a complexed autophagy. Finally, our study revealed that the combination of GNS561 and remdesivir was associated with a strong synergistic antiviral effect against SARS-CoV-2. Overall, our study highlights GNS561 as a powerful drug in SARS-CoV-2 infection and supports that the hypothesis that autophagy inhibitors could be an alternative strategy for SARS-CoV-2 infection.

## Introduction

Since December 2019, severe acute respiratory syndrome coronavirus 2 (SARS-CoV-2/2019-nCoV, also called **<u>co</u>**rona**<u>vi</u>**rus **<u>d</u>**isease of 2019, Covid-19) has spread quickly worldwide and has qualified as a pandemic *(1)*. As of October 2020, more than 29 million cases and 927,000 deaths have been reported worldwide (World Health Organization data). SARS-CoV-2 infection leads to a range of symptoms from mild fever to acute respiratory distress syndrome classifying this disease into several clinical categories, including asymptomatic, mild, moderate, severe and critical infection *(2)*.

Thus, there is an urgent need for a safe and powerful, accessible and affordable drug to prevent and control early disease, restrain, and slow the epidemic. To date, researchers have reported at least 87 drugs as potential Covid-19 therapies. Among them, 68 were tested *in vitro* and several are in clinical testing or under preclinical development after receiving FDA (Food and Drug Administration) approval *(3)*. To date, 3,328 clinical trials concerning Covid-19 disease are underway worldwide (www.clinicaltrials.gov). Among the investigated molecules, most are therapeutic repositioning compounds such as chloroquine (CQ), an older FDA-approved anti-malaria drug, baricitinib, dexamethasone, or the experimental anti-Ebola virus drug remdesivir *(4)* that has recently been given emergency approval as a Covid-19 treatment.

The diversion of the autophagy mechanism by viruses, including coronaviruses, allows for viral replication *(5)*. More specifically, coronaviruses were found to use double membrane vesicles to enhance the efficiency of virus replication *(6)*. Additionally, it has been suggested that coronaviruses induce autophagy *(7)*, although the mechanism of SARS-CoV-2 remains to be determined. During the Covid-19 global pandemic, researches turned to autophagy inhibitors as a target mechanism effective in the inhibition of SARS-CoV-2 infection. CQ and hydroxychloroquine (HCQ), a derivative of CQ, are autophagy inhibitors and efficient against a large variety of pathologies, including cancer, malaria and systemic lupus erythematous, and they have also been shown to have antiviral effects *in vitro (8)*. Both were reported to have an effect on lysosomes by inhibiting the pH-dependent replication steps of several viruses, including flaviviruses, retroviruses, and coronaviruses, including SARS-CoV-2 *(9)*. It turned out that the autophagy mechanism acts as an antiviral defense in the early step of viral infection *(5)*.

Our team developed an autophagy inhibitor, GNS561, which is a small basic lipophilic molecule that induces lysosomal dysregulation, as proven by the inhibition of late-stage autophagy *(10)*. GNS561 is currently in development for oncology indications (NCT03316222) and it is being tested in a clinical trial of patients infected by Covid-19 (NCT04333914). The objectives of the current study were i) to evaluate the *in vitro* antiviral activity of GNS561 against SARS-CoV-2, ii) to assess the mechanism of blocking the viral replication by inhibition of autophagy, and iii) to demonstrate a synergistic combination with remdesivir.

## Materials and Methods

### Cell culture and virus

Vero E6 cells (African green monkey kidney, American Type Culture Collection ATCC® CRL-1586™) were cultured using Minimum Essential Media (MEM) (Life Technologies, Carlsbad, CA, USA) supplemented with 10% fetal bovine serum (FBS) (HyClone, Logan, UT, USA) and 1% L-glutamine (Life Technologies). Cells were maintained at 37°C in the presence of 5% CO_2_ and 95% air in a humidified incubator.

SARS-CoV-2 strains, IHU-MI3 -MI6 and USA-WA1/2020 were provided by the IHU Mediterranean Infection and BEI Resources (Manassas, VA), respectively. Both viral strains were propagated in the Vero E6 cell line and titrated by the TCID_50_ method followed by storage of aliquots at −80°C until further use in the experiments.

### In vitro cytotoxicity assay

Cell viability was assessed using the CellTiter-Glo® Luminescent Cell Viability Assay following the manufacturer’s protocol (Promega, Madison, WI, USA). The luminescence was recorded using an Infinite F200 Pro plate reader (Tecan, Männedorf, Switzerland). Cytotoxicity was expressed as the concentration of test compounds that inhibited cell proliferation by 50% (CC_50_) and calculated using the Chou and Talalay method *(11)*. Every analysis was performed in technical triplicate, and three independent experiments were performed.

### Antiviral Activity Assay

A confluent monolayer of Vero E6 cells prepared in a 96-wells cell culture microplate was followed by treatment either with GNS561, CQ, remdesivir (0.1-50 µM for GNS561 and remdesivir, 0-200 µM for CQ), or vehicle control for 2 hours. Treated cells were then infected with SARS-CoV-2 (USA-WA1/2020 and IHU-MI3 -MI6) at an MOI of 0.1 for 24h. The yield of progeny virus production was assessed on supernatant of infected cells using a specific qRT-PCR. Briefly, a one-step qRT-PCR was conducted in a final volume of 25 μL containing 5µL of extracted viral RNA from the SuperScript™ III Platinium™ One-Step qRT-PCR Kit (Life Technologies), 20 µL of a mix containing the probe/primer mix (**Table 1**), and Super Script master mix (Life Technologies). Quantitative PCR measurement was performed using the Cobas z 480 PCR system (Roche, Bâle, Switzerland). Data were analyzed with the LightCycler 480 SW 1.5 software (Roche) according to the manufacturer’s protocol. Fold changes were calculated as the relative expression of the E gene after normalizing to both infected and untreated cells. A melting curve analysis was performed after amplification to verify the accuracy of the amplicon. Every analysis was performed in technical triplicate, and three independent experiments were performed.

**Table 1.**
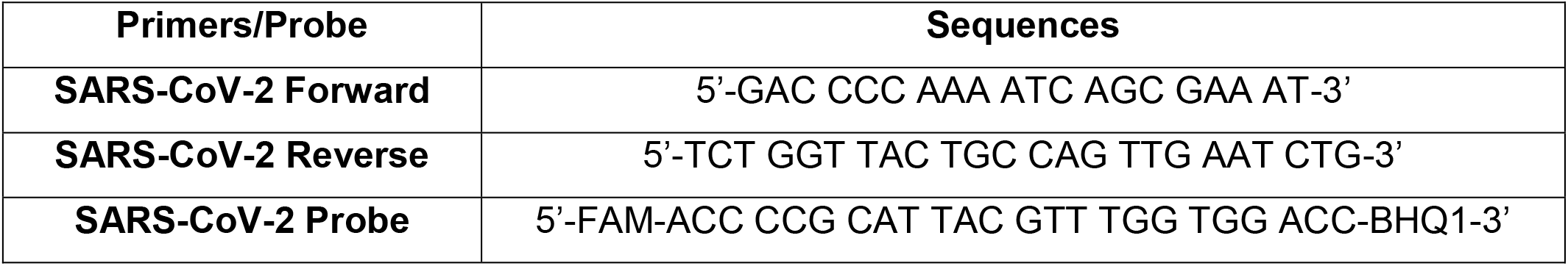
Primers and probe sequences for SARS-CoV-2 q-RTPCR investigation.

### Cell imaging

The autophagy flux of the cells was investigated in infected or non-infected Vero E6 cells by two different techniques. First, the autophagy was investigated by immunofluorescence. Vero E6 cells cultured on coverslips were initially treated or not by a fluorescent analog of GNS561, GNS561G, for 2 hours and were then infected by SARS-CoV-2 (IHU-MI6, MOI 0.1). After that, cultured cells were rinsed with PBS and fixed with 3% paraformaldehyde for 20 min at 4°C. Cells were washed again with PBS and blocked with 1× PBS supplemented with 5% FBS (blocking buffer) for 25 min at room temperature (RT), followed by a permeabilization step with 3% Triton™ X-100 (Sigma Aldrich) for an additional 5 min in the blocking buffer. Then, the coverslips were incubated in blocking buffer for 45 min at room temperature with antibodies against LC3-II (Sigma Aldrich, 1:200), LAMP2 (DSHB, 1:250), and SARS Spike protein (Thermo Fisher, 1:250), and with Phalloïdin-647 (Life Technologies, 1:250) and 4’,6-diamidino-2-phenylindole (DAPI, Life Technologies, 1:250) to reveal actin and the nucleus, respectively. The coverslips were then washed three times with PBS and incubated for 30 min at RT in 1:1000 diluted secondary antibodies. The secondary antibodies used to reveal LC3 and LAMP2 were Alexa 594-labeled anti-rabbit secondary antibody and Alexa 546-labeled anti-mouse secondary antibody (Invitrogen, USA), respectively. Once again, the coverslips were washed three times with PBS and then mounted using Mowiol (Sigma Aldrich, USA) overnight at 4°C. Fluorescence images were acquired using an LSM 800 Airyscan confocal microscope (Zeiss, Germany) with a 63× oil objective. Six randomly selected microscopic fields of cells were acquired and collected by Zen 3.0 (Blue Edition) software (Zeiss).

Autophagy was then investigated by electron microscopy. Vero E6 cells were fixed for at least 1 h with glutaraldehyde 2.5% in 0.1 M sodium cacodylate buffer. For resin embedding, the cells were washed three times with a mixture of 0.2 M saccharose/0.1 M sodium cacodylate. Cells were then postfixed for 1 h with 1% OsO4 diluted in 0.2 M potassium hexa-cyanoferrate (III) / 0.1 M sodium cacodylate solution. After three 10 min washes with distilled water, the cells were gradually dehydrated with ethanol by successive 10 min baths in 30, 50, 70, 96, 100, and 100% ethanol. Substitution was achieved by successively placing the cells in 25, 50, and 75% Epon solutions for 15 min each. The Vero E6 cells were placed for 1 h in 100% Epon solution and in fresh Epon 100% overnight under vacuum at room temperature. Polymerization occurred with the cells in 100% fresh Epon for 72 hours at 60°C. All solutions used above were 0.2 µm filtered. Ultrathin 70 nm sections were cut using a UC7 ultramicrotome (Leica) and placed on HR25 300 Mesh Copper/Rhodium grids (TAAB). Sections were contrasted according to the methods of Reynolds *(12)*. Electron micrographs were obtained on a Morgagni 268D (Philips / FEI) transmission electron microscope operated at 80 keV TEM.

### Western blot assay

The light chain 3-phosphatidylethanolamine-conjugate (LC3)-II (Sigma Aldrich, 1:3000) and glyceraldehyde-3-phosphate dehydrogenase (GAPDH) (Abnova, Taipei, Neihu, Taïwan, 1:5000) protein expression levels were investigated by western blotting using Imager iBrightCL1000 (Thermo Fisher). Two hours before Covid-19 infection (IHU-MI3 strain, MOI 0.25), cells were treated with 25 µL of the investigated drugs and then infected with 50 µL of virus suspension for an extra 24 hours at 37°C in the presence of 5% CO_2_ and 95% air in a humidified incubator. During the last 2 hoursours, cells were treated or not with 100 nM bafilomycin A1 (Baf-A1, Sigma Aldrich), an autophagy flux inhibitor.

### Combination assay

To evaluate the synergetic effect of the investigated drugs, 25,000 cells/well were seeded into 96-well plates, cultured overnight and then treated in triplicate with GNS561 or remdesivir, either alone or in various combinations, for 26 h. Two hours after drug treatment, the cells were infected with the SARS-CoV-2 IHU MI6 strain for 24 additional hours. At the end-point, antiviral effects were investigated on supernatant of infected cells using CellTiter-Glo® and qPCR as described below. Data were analyzed using the MacSynergy™ II Excel Spreadsheet based on the Bliss Independence model as previously described *(13)*. Briefly, this program compares the additive *versus* synergism effect to the experimental combined-drug effect. This can be described by a surface plot of peaks and valleys of synergy and antagonism values interpreted in log volumes (LV) informing a quantitative measure of synergy or antagonism over the sum of the concentrations tested. LV ≥20, −20< LV <20 and LV ≤-20 indicate synergy, additive effect and antagonism, respectively.

## Results

### GNS561 exhibits strong antiviral activity against SARS-CoV-2

As illustrated in **Fig. 1** and **Table 2**, GNS561 showed the most potent antiviral effect against SARS-CoV-2 compared to CQ and remdesivir. Indeed, GNS561 exhibited a 0.006 µM median effective concentration (EC_50_) with 2.0 µM cytotoxicity concentration (CC50), whereas CQ and remdesivir presented 0.10 µM EC_50_ (73.2 µM CC50) and 1.2 µM EC_50_ (> 100 µM CC50), respectively. More precisely, the GNS561 inhibitory concentration was 16 and 200 times higher than CQ and remdesivir, respectively. Moreover, we confirmed the strong antiviral effect of GNS561 using another SARS-CoV-2 strain (IHU-MI6). As illustrated in **Table 2**, GNS561 presented a 0.03 EC_50_ and 0.07 μM EC_90_. To summarize, our data showed that GNS561 activity is higher compared to CQ and remdesivir respectively.

**Table 2:**
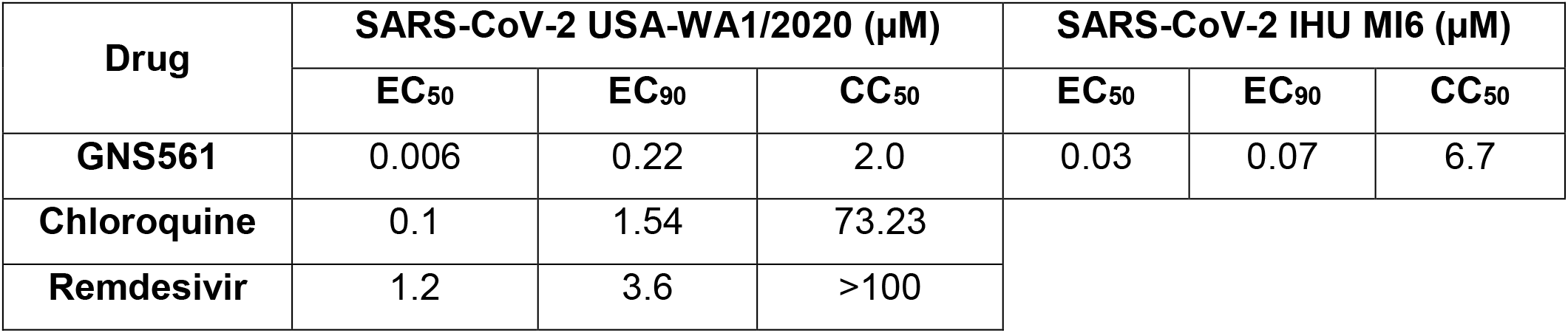
Antiviral activity and cytotoxicity of GNS561, chloroquine and remdesivir against SARS-CoV-2 in Vero E6 cells. After 2 hours of treatment with different doses, Vero E6 cells were infected with SARS-CoV-2 virus IHU-MI6 or USA-WA1/2020 strains (MOI of 0.1) for 24 hours. The inhibitory concentration (IC) and cytotoxicity (CC) in µM were investigated using qRT-PCR and viability assays, respectively.

| Drug | SARS-CoV-2 USA-WA1/2020 ($\mu\text{M}$) | | | SARS-CoV-2 IHU MI6 ($\mu\text{M}$) | | |
| --- | --- | --- | --- | --- | --- | --- |
|  | EC<sub>50</sub> | EC<sub>90</sub> | CC<sub>50</sub> | EC<sub>50</sub> | EC<sub>90</sub> | CC<sub>50</sub> |
| **GNS561** | 0.006 | 0.22 | 2.0 | 0.03 | 0.07 | 6.7 |
| **Chloroquine** | 0.1 | 1.54 | 73.23 |  |  |  |
| **Remdesivir** | 1.2 | 3.6 | >100 |  |  |  |

**Figure 1.**
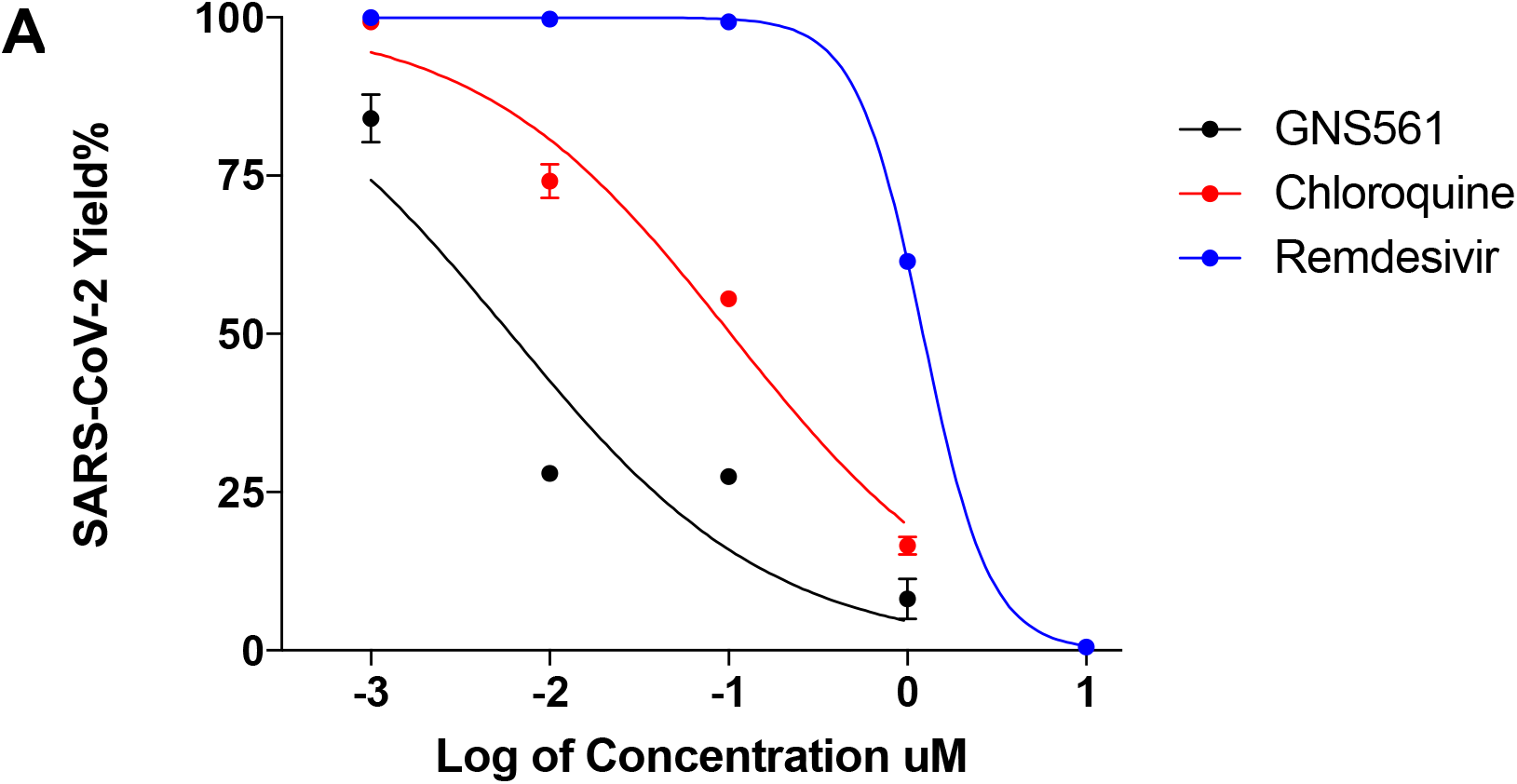
The *in vitro* antiviral activities of the tested drugs against SARS-CoV-2. After 2 hours of treatment with different doses, Vero E6 cells were infected with USA-WA1/2020 strain. The viral yield was quantified by qRT-PCR. The left axe of the graph represent the percentage of SARS-CoV-2 yield. The results obtained for GNS561, CQ and remdesivir treatments are represented in black, red and blue, respectively. Values are means of at least duplicate assays conducted in triplicate.

### Antiviral activity of GNS561 is mediated by autophagy inhibition

First, we investigated SARS-CoV-2 repartition in the intracellular compartment of the Vero E6 cells using electron microscopy. We were able to distinguish (1) the entry of SARS-CoV-2 at the periphery of Vero E6 cells (red arrow), (2) the presence of endocytic vesicles in the cytoplasm with clathrin-coated vesicles (yellow arrow) and (3) the virus-producing cell with vacuoles filled with nascent particles (red asterisk) (**Fig. 2A**). GNS561 treatment leads to an accumulation and an increase in the volume of autophagic vacuoles in the cytoplasm with the presence of multilamellar bodies characteristic of a complexed autophagy (**Fig. 2A**).

**Figure 2.**
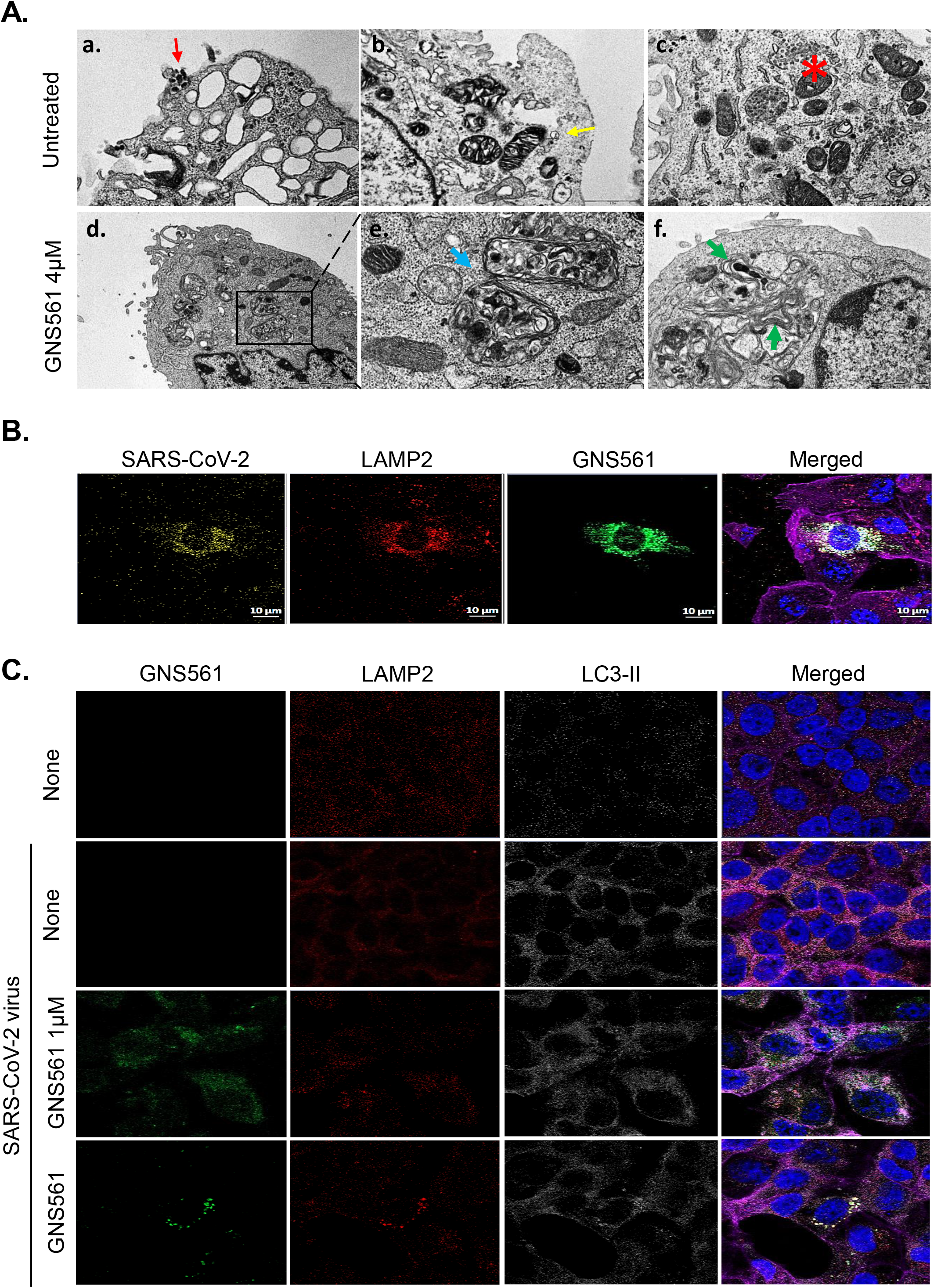
GNS561 leads to autophagy blockage. After 2 hours of treatment with different doses, Vero E6 cells were infected with SARS-CoV-2 IHU-MI6 strain. (**A**) Electron microscopy pictures illustrating infected cells without (upper panel) or with GNS5561 treatment (lower panel). The presence of the virus is indicated using a red arrow, endocytic vesicles in the cytoplasm with clathrin-coated vesicles with yellow arrows, autophagy vacuole with blue arrows and multilamellar bodies with green arrows. Vacuoles filled with nascent particles are illustrated using a red asterisk. (**B**) Confocal images showing the localization of SARS-CoV-2 (yellow) inside LAMP2-positive lysosomes (red) together with GNS561 (green). (**C**) Confocal pictures illustrating GNS561 (green), LC3 (white) and LAMP2 (red) expression in Vero E6 cells with nuclei in blue and actin in purple.

Based on these results, we next investigated the location of SARS-CoV-2 in the presence of our antiviral compound in the cellular compartments associated with the autophagy mechanism. As depicted in **Fig. 2B**, we noted that SARS-CoV-2 was located in LAMP2-positive lysosomes similar to GNS561. Next, we showed that SARS-CoV-2 infection leads to an increase in size of LAMP2 particles associated with a reorganization of intracellular LC3-II expression, also called spots (**Fig. 2C**). In a dose-dependent manner, the size of LC3-II spots increased following GNS561 treatment, suggesting that the way of autophagy is impacted.

Finally, LC3-II expression was investigated by western blotting after treatment of Vero E6 cells with 100 nM bafilomycin A1 (Baf A1, an autophagy flux inhibitor) during the last 2 hours of the experiment. As depicted in **Fig. 3A**, LC3-II was increased following SARS-CoV-2 infection, suggesting that the infection of cells was specifically mediated by autophagic flux, as no differences were observed following BafA1 treatment as a control. GNS561 treatment led to an increased in LC3-II protein expression in a dose-dependent manner, suggesting effective modulation of the autophagy. Moreover, the accumulation of LC3-II protein in GNS561-treated cells was not enhanced in the presence of Baf-A1, confirming the specificity of GNS561 to inhibit autophagy (**Fig. 3B**). Interestingly, normalized LC3-II protein expression was increased in SARS-CoV-2-infected cells treated with GNS561 and decreased after Baf-A1 treatment, suggesting that the compound altered autophagosome synthesis (**Fig. 3C**). Overall, our results indicate that GNS561 impacts on autophagy mechanism and subsequent inhibition of viral multiplication.

**Figure 3.**
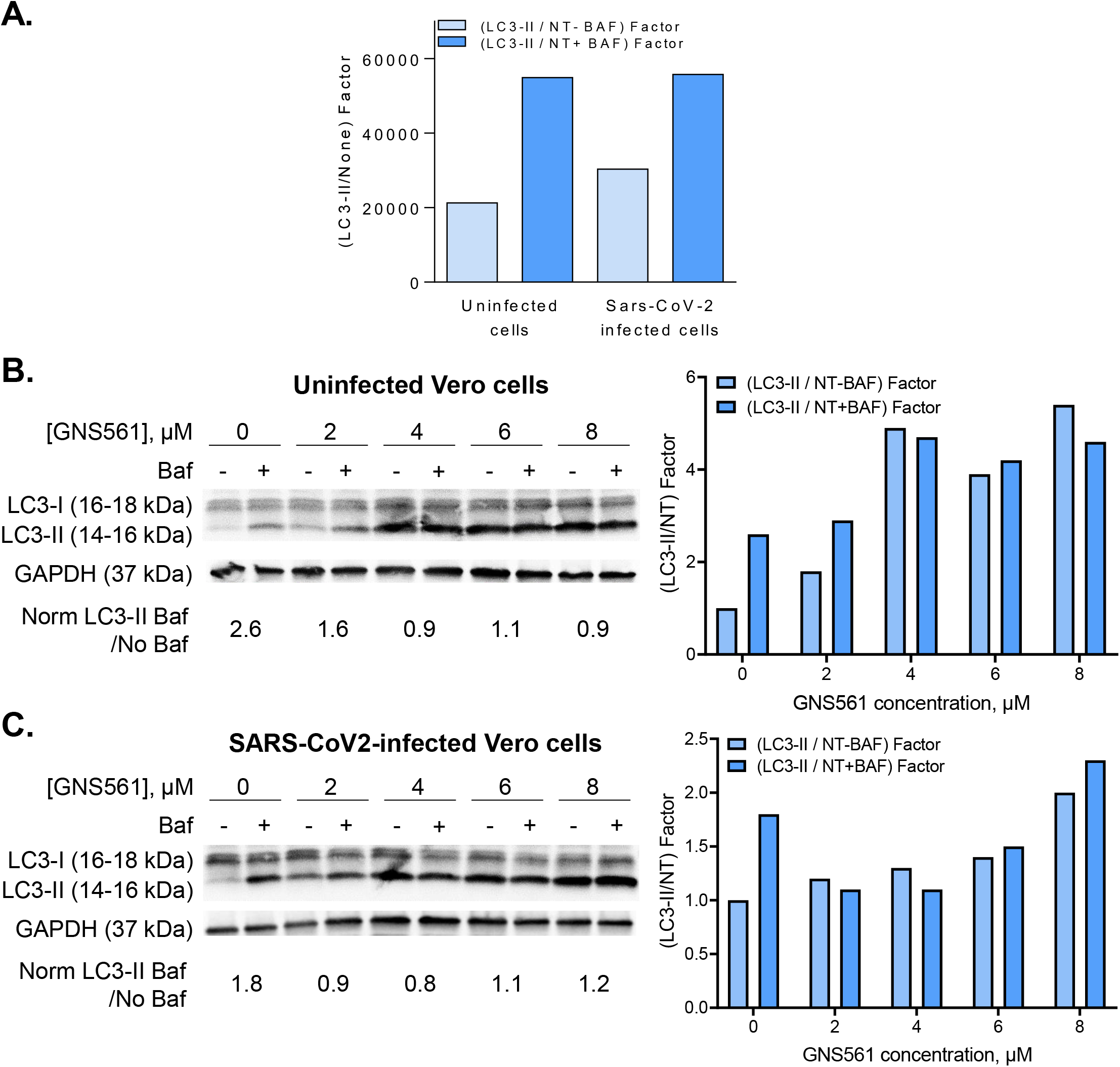
Antiviral activity of GNS561 is mediated by the autophagy inhibition. LC3-I and LC3-II protein expression was evaluated by western blotting and normalized for LC3-II protein for (**A**) cells infected or not infected by SARS-CoV-2. Autophagy inhibition was evaluated in the presence of absence of bafilomycin A1 (Baf-A1) treatment added 2 hours before the end of the experiment. Cells were treated with GNS561 (2, 4, 6 and 8 µM) in (**B**) uninfected or (**C**) SARS-CoV-2 infected (MOI of 0.25) conditions. Experiments were performed in triplicate.

### GNS561 and remdesivir combination presents a powerful synergic effect against SARS-CoV-2 infection *in vitro*

It has been proposed that a combination of drugs with independent mechanisms of action could be an effective strategy in SARS-CoV-2 infection. To test this, Vero E6 cells were treated with different concentrations of GNS561 or remdesivir either alone or in combination, with drug synergy/antagonism assessed by the MacSynergy method. As illustrated in **Fig. 4**, our results showed a strong synergistic effect on the antiviral activity of GNS561 at all tested concentrations with remdesivir in the range of 0.1 to 0.5 μM.

**Figure 4.**
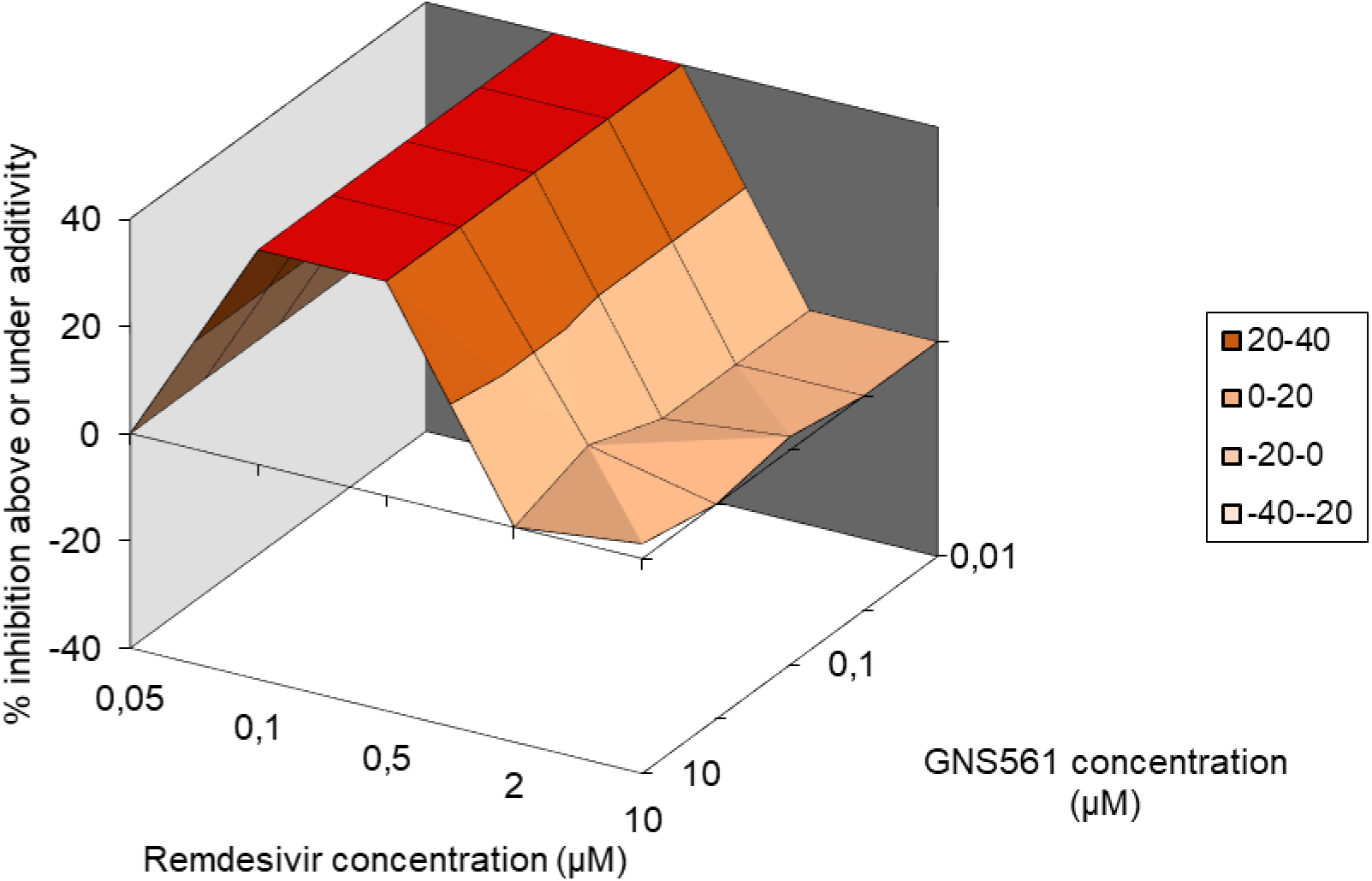
Synergistic antiviral action of GNS561 with remdesivir. After 2 h of treatment with different doses of GNS561 or remdesivir either alone or in combination, Vero E6 cells were infected with SARS-CoV-2 IHU-MI6 strain (MOI of 0.1). Synergy graph generated using MacSynergy illustrating the antiviral effect of GNS561 in combination with remdesivir.

## Discussion

In this work, we have demonstrated that, GNS561 exhibits a much stronger antiviral effect against SARS-CoV-2 compared to CQ and remdesivir. Liu. J et al. reported that CQ and remdesivir efficiently inhibited SARS-CoV-2 infection *in vitro (14)*. Indeed, remdesivir was found to be effective against SARS-CoV-2 with an EC_50_ of 0.77 *(14, 15)*. Moreover, CQ and HCQ have been proposed as treatment for SARS-CoV-2 infection *(14, 16, 17)*. At every tested MOI (0.01, 0.02, 0.2, and 0.8), the EC_50_ for CQ (2.71, 3.81, 7.14 and 7.36 μM) was lower than that of HCQ (4.51, 4.06, 17.31 and 12.96 μM). Interestingly, the EC_50_ values of CQ that they obtained are barely higher than their previous work (from 1.13 μM to 7.36 μM). It appears that they may have underscored the adaptation of the virus in cell culture that substantially increased its viral infectivity upon continuous passaging. Indeed, GNS561 exhibits one of the most powerful antiviral effect *in vitro* described in the literature so far against SARS-CoV-2.

Our study showed that i) SARS-CoV-2 was located in LAMP2-positive lysosomes similar to GNS561. ii) SARS-CoV-2 infection led to an increase in LAMP2 particles associated with an autophagy induction. Some viruses hijack the autophagy mechanism to allow their entry and replication, as previously reported for SARS-CoV replication *(18–22)*. CQ and HCQ, two autophagy inhibitors, were reported to elicit antiviral activity through the alteration of endosomal pH and the inhibition of lysosomal fusion to viral endosomes, leading to the reduction of viral replication and the protection of cells from SARS-CoV-2-induced cell death *(7)*. Here, we proposed GNS561, a late-stage autophagy inhibitor, as an effective antiviral compound in the inhibition of autophagy during SARS-CoV-2 infection targeting autophagy vacuoles during the early steps in the viral replication cycle. Indeed, this is the first observation of autophagy vacuole blockade during SARS-CoV-2 infection that shows characteristics of complexed autophagy. Moreover, many mechanisms have been described to target the viral replication cycle associated with lysosomotropic agent treatment have been proposed *(23–25)*. Further investigations are needed to dissect the different target mechanisms of the autophagy pathway of the GNS561 drug in SARS-CoV-2 infection.

Today, direct active antiviral drugs are the dominant class of antiviral drugs in use. They are highly successful against clinically important infections such as human immunodeficiency viruses, hepatitis C virus and several others. Remdesivir, a nucleotide analog, recently showed significant clinical results in the treatment of Covid-19 patients *(26, 27)*. Nonetheless, obstacles such as drug-resistant viruses or emerging viruses with no available selective antivirals in the market call for the utilization of an armament of simple compounds that target host factors and show broad-spectrum antiviral activities. Thus, there is some interest in using drug combinations against SARS-CoV-2. Interestingly, it was recently reported that nitazoxanide in combination with amodiaquine, umifenovir or remdesivir presented a significant synergy against SARS-CoV-2 *(28)*. It was also reported that remdesivir and HCQ exhibited a strong antagonism *(28)*. Here, we showed that a combination of the replication inhibitor, remdesivir with the autophagy inhibitor, GNS561 demonstrated strong synergy *in vitro*, supporting future clinical evaluation of such combination regimens. Overall, the importance of both drug repurposing and preclinical testing of drug combinations must be taken into consideration for potential therapeutic use against SARS-CoV-2 infection.

There are conflicting data regarding the efficacy and toxicity of HCQ alone or in combination for the treatment of Covid-19-positive patients *(29)*. Of concern are the side effects were reported concerning CQ treatment. Indeed, although CQ and HCQ present safety profiles in clinical use, side effects have been reported, including gastrointestinal upset *(30)*, retinal toxicity *(31, 32)*, cardiomyopathy *(33)* and heart rhythm disturbances *(34)*. Compared to HCQ and CQ, GNS561 presents several advantages, including no phototoxicity and no electrocardiogram modification with a markedly more potent activity the SARS-CoV-2 through its anti-autophagy mechanism. Interestingly, GNS561 is currently being tested in moderate Covid-19 patients in a Phase II clinical study. Due to this original mechanism of action, the focus of the Covid-19 patients is located is the early phase before the cytokine storm.

Taken together, due to the high potency of GNS561 against Covid-19 and its good safety profile observed in phase II clinical trials, GNS561 constitutes a promising treatment for Covid-19. In addition, considering the continuous spread of major viral pathogens as well as unpredictable viral outbreaks of old or novel coronaviruses, it seems advisable to have an arsenal of countermeasures ready for the prevention of global health crises.

## Acknowledgments

This work was supported in part by the funding from the Emory Center for AIDS Research (5P30-AI-50409 to RFS).

## Author contributions

PH, EB, SM and RFS conceived and designed experiments. EB, KZ, JA, JPB and SM realized experiments. PH, BLS, JLM, SM and RFS supervised the study. PH, EB, SM and RFS wrote the manuscript.

